# Evolutionary rescue through partly heritable phenotypic variability

**DOI:** 10.1101/092718

**Authors:** Oana Carja, Joshua B. Plotkin

## Abstract

Environmental variation is commonplace, but unpredictable. Populations that encounter a deleterious environment can sometimes avoid extinction by rapid evolutionary adaptation. Phenotypic variability, whereby a single genotype can express multiple different phenotypes, might play an important role in rescuing such populations from extinction. This type of evolutionary bet-hedging need not confer a direct benefit to a single individual, but it may increase the chance of long-term survival of a lineage. Here we develop a population-genetic model to explore how partly heritable phenotypic variability influences the probability of evolutionary rescue and the mean duration of population persistence in changing environments. We find that the probability of population persistence depends non-monotonically on the degree of phenotypic heritability between generations: some heritability can help avert extinction, but too much heritability removes any benefit of phenotypic variability. We discuss the implications of these results in the context of therapies designed to eradicate populations of pathogens or aberrant cellular lineages.

## Introduction

The very first study in experimental evolution, led by W. D. Dallinger in the 1880s, attempted to demonstrate that populations can rapidly adapt to environmental change and that evolutionary rescue of a population from extinction depends on the rate of change (Dallinger, 1887). Evolutionary rescue is the process by which a population is able to recover from abrupt environmental changes that would otherwise lead to a demographic decline and eventual extinction (Gomulkiewicz and Holt, 1995; Bell and Gonzalez, 2009; Alexander et al., 2014; Gonzalez et al., 2013; Lindsey et al., 2013; Carlson et al., 2014). Evolutionary responses that allow populations to adapt to change on a sufficiently fast time scale to prevent extinction have been the focus of considerable experimental and theoretical interest, across diverse biological systems. In the field of conservation biology, questions of rescue are framed around ensuring the survival of species in deteriorating global habitats (Palumbi, 2001; Gonzalez et al., 2013; Davis et al., 2005). In contrast, in clinical contexts, the goal is eradication of pathogens or harmful populations of cells (Gonzalez et al., 2013; Alexander et al., 2014). These two bodies of work share a common thread (as Maynard Smith (1989) emphasized, adaptation in threatened populations is not like ordinary adaptation, it is a race against extinction), yet each presents unique diffculties.

Here we focus on the questions of population eradication that arise in medically-relevant settings, where populations are often surprisingly resilient or recalcitrant to treatment due to evolutionary adaptation. In particular, we ask: how does heritable phenotypic variability alter the probability of evolutionary rescue? We study the evolutionary advantage of heritable phenotypic variability in populations of non-constant size. We determine the probability of rescue and the mean time to extinction in changing environments, through both analytical approximation and Monte Carlo simulations of population genetic models.

We are motivated in asking this question by well-documented examples of phenotypic heterogeneity used as evolutionary bet-hedging strategy in volatile environments. Classic examples include the bifurcation of a genotypically monomorphic population into two phenotypically distinct bistable subpopulations (Dubnau and Losick, 2006); bacterial persistence (Lewis, 2010; Cohen et al., 2013; Kussell et al., 2005; Sharma et al., 2015), whereby a genetically identical bacterial population survives periods of antibiotic stress by producing phenotypically heterogeneous sub-populations, some of which are drug-insensitive (Lewis, 2007); or quiescent phenotypes in cancer cell populations (Aguirre-Ghiso, 2007; Sharma et al., 2010; Smith et al., 2016), which are transient phenotypic (epi)states protected from the action of drugs. These dormant phenotypic states confer the population with some degree of phenotypic heterogeneity, helping it withstand periods of environmental stress. Phenotypes may be partially heritable upon cellular division, so that the offspring cell can sometimes “remember” and express the phenotypic state of its parent, or sometimes switch between phenotypic states at rates that greatly exceed those of genetic mutations (Balaban et al., 2004; Van den Bergh et al., 2016). Partial phenotypic inheritance through epigenetic mechanisms can lead to faster rates of adaptation and environmental tracking than genetic mutations alone. Even though persisters in such populations rely on a non-genetic form of inheritance, the rate of ‘phenotypic mutation’ is itself likely under genetic control (Levin and Rozen, 2006).

Epigenetic bet-hedging strategies that use dynamic regulation of phenotypic variability can allow a population to persist and escape extinction, until more permanent genetic strategies arise. Many studies have addressed questions of genetic responses in evolutionary rescue (Lindsey et al., 2013; Uecker and Hermisson, 2015; Wilson et al., 2017; Uecker et al., 2014; Orr and Unckless, 2008, 2014; Alexander et al., 2014; Martin et al., 2013). Less attention has been given to the potential impact of phenotypic variability and of stochastic switching on evolutionary rescue (Ashander et al., 2016; Chevin et al., 2013). A recent study integrated stochastic demography with quantitative genetic theory in a model with evolving plasticity to explore the probability of rescue from a single, abrupt change in the environmental conditions (Ashander et al., 2016). Evolving plasticity was shown to facilitate evolutionary rescue unless the new environment is unpredictable.

Epigenetic plasticity as studied in Ashander et al. (2016) can cause phenotypes to differ widely within a lineage, whereas purely genetically encoded phenotypes only allow offspring phenotypically similar to the parents. The type of phenotypic variability we explore here can produce phenotypic heterogeneity with familial correlations intermediate to these two extremes – for example, as observed in the contributions of DNA methylation variation to the heritability of phenotypes in Arabidopsis thaliana (Johannes et al., 2009) (Carja et al., 2014a; Carja and Plotkin, 2017). This type of partly heritable phenotype is commonplace in medical settings, and its role in evolutionary rescue is yet to be understood.

We explore the evolutionary fate of a population that experiences either one sudden shift in the environmental regime, or many periodic changes in the environment. In the case of a single abrupt environmental change, we study the probability of rescue when one mutant allele permits the expression of multiple phenotypic states. We imagine these phenotypic states as partially heritable, so that the phenotype expressed by an individual will be inherited by the offspring with some probability, *p*. We call this probability p the phenotypic memory. We are especially interested in the long-term fate of the population as a function of the variance in expressible phenotypes that the mutant allele confers, and also as a function of the amount of phenotypic memory between generations.

Our paper starts by specifying a mathematical model, based on birth-death processes, for populations subject to environmental change and to the introduction of a mutant allele that permits a range of expressible, partly heritable phenotypes. We pose our research question in terms of analyzing the long-term probability of extinction of such a population. We show that after one abrupt environmental change, the probability of evolutionary rescue is significantly increased when a new phenotypically variable allele is introduced in the population, and this increase critically depends on the phenotypic memory of individuals expressing the variable allele. When the population experiences multiple environmental changes, the mean time to population extinction also increases for phenotypically heterogeneous populations (i.e. population persistence increases) and this increase depends non-monotonically on the phenotypic memory of the mutant allele, *p*. We provide a simple intuition for the complex dependence of evolutionary rescue on the degree of phenotypic memory, and we discuss the implications of our results for the eradication of evolving populations in medical contexts.

### Model

We use a continuous-time birth-death model to describe changes in allele numbers in a finite population of changing size *N*, with carrying capacity *K*. Each individual’s genotype is defined by a single biallelic locus *A*/*a*, which controls its phenotype. The *A* allele encodes a fixed phenotypic value, whereas individuals with the *a* allele may express a wider range of phenotypes, drawn from a fixed distribution. Here we analytically study the case where the *a* allele has access to two different phenotypic values. By simulation we also explore cases where the *a* allele has access to more than two phenotypic states. The phenotypic values available to the a allele are chosen such that both alleles *A* and a have the same expected fitness, so that the only difference between them is the possibility of (partly heritable) phenotypic variability. We analyze the probability of rescue as a function of the phenotypic variance and the phenotypic memory associated with the a allele.

We study two sets of questions related to phenotypic variability and persistence. In the first set of questions, the population, assumed to be initially fixed for the wild-type allele A, experiences a single abrupt change in the environmental regime at time *t* = 0. This environmental shift is expected to lead to a demographic decline in the population, meaning that death rates exceed birth rates for allele *A*. We ask what is the probability of rescue when there is a positive mutation rate to the phenotypically variable allele *a*, which can provide an adaptive benefit. In this situation there is a race between extinction of the population, and the origin and establishment of a phenotypically variable *a*-lineage. We also ask what is the probability of evolutionary rescue from standing genetic variation – that is, if the a allele is already present in the population at some frequency at time *t* = 0. We study how the probability of rescue depends upon parameters such as the population size at time *t* = 0, the phenotypic variance of the *a* allele, the phenotypic memory of the a allele, and either the mutation rate towards *a* or the initial frequency of *a*.

In the second set of studies, we assume a population otherwise fixed on the non-variable *A* allele with one phenotypically-variable *a* allele introduced at time *t* = 0; but here we assume multiple epochs of environmental changes, occurring periodically. The question of persistence is framed in terms of the mean time to extinction of the population, as a function of the environmental period, the phenotypic variance and the phenotypic memory available to the a allele. In this case, the mapping from phenotype to fitness depends on the environmental regime, and it is chosen so that both alleles have the same expected fitness across environments.

### Evolutionary rescue from a single environmental change

The constant-environment model studies how a novel allele that can express multiple phenotypes might alter the probability of evolutionary rescue of a population otherwise headed towards extinction. We describe the population using a continuous time birth-death model. We assume the death rates are density independent and we study both density-dependent and density-independent birth rates. Individuals of the wild type *A* and mutant type *a* each give birth and die according to the following per-capita rates:

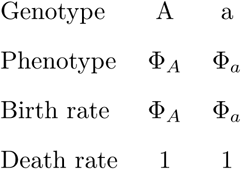

for the density-independent model and

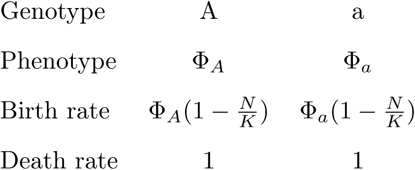

for the density-dependent model. Here Φ_*A*_ is a fixed number and Φ_*a*_ is a discrete random variable with probability mass on two values, Φ_*a-*_ and Φ_*a*+_. N denotes the current population size, and K the carrying capacity. The death rates of the two alleles are both assumed to be unity. The values Φ_*A*_ and Φ_*a*_ are constrained so that the two alleles have the same mean fitness: Φ_*A*_ = 𝔼(Φ_*a*_). We study fitness functions with equal means so that we can focus our analysis on the effect of variance in phenotypes expressed by allele a, Var(Φ_*a*_), and not on any mean-fitness effects. An illustration of this model is presented in **Figure 1A**. In our analysis of this model we initiate the population in a regime where the wild-type allele has a higher per-capita death rate than its maximum possible birth rate, so that a wild-type population is expected to go extinct fairly quickly. We analyze the conditions under which the mutant allele a (arising either by mutation or from standing variation at time *t* = 0) will rescue the populations from extinction. We compare this analysis with Monte Carlo simulations in which we record the proportion of replicate simulations in which rescue occurs.

### Population persistence in periodically changing environments

In our analysis of periodic environmental changes we assume that the population experiences two different types of environments, *E*_1_ and *E*_2_, which alternate deterministically every *n* generations, so that both environments are experienced every 2*n* generations. We assume that one environment is more favorable to one allele, and the other environment to the other allele. We study a scenario where the phenotypically-variable allele *a* has lower expected fitness than the wild-type allele in one of the environmental regimes; and higher expected fitness than the wild-type in the other regime. In the case of persister phenotypes in bacteria, for example, the environment that is detrimental to the allele a corresponds to the “no antibiotic” regime; whereas the persister a has a higher expected fitness than wild-type in the presence of antibiotic pressure.

We choose phenotypic ranges and fitness functions so that the mean fitness expressed by each of the two genotypes are equal, averaging over the two environmental regimes. This setup allows us to focus on the evolutionary advantage of phenotypic variance of *a*, Var(Φ_*a*_), and to study its consequences for population persistence without conflating its effect with any mean-fitness advantage of the a allele over the A allele.

Individuals of the wild type A and mutant type a each give birth and die according to the following per-capita rates:

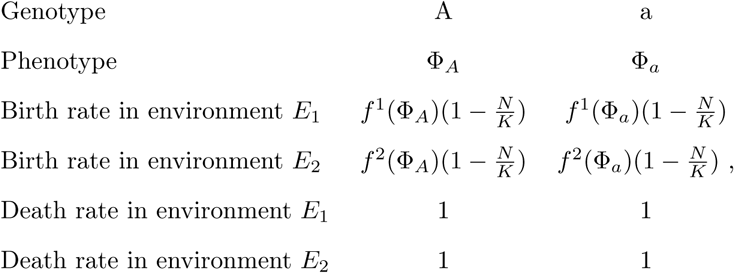

where Φ_*A*_ and Φ_*a*_ denote random variables, although Φ_*A*_ is in fact deterministic with zero variance. The functions *f*^*i*^: ℝ *→* ℝ (*i* ∊ {1, 2}) map phenotype to fitness in each of the two environments, and *f*^1^ is taken to be the identity function. We assume that both alleles have the same mean fitness in their preferred environment, and the same mean fitness in their unpreferred environment: 𝔼(*f*^1^(Φ_*A*_)) = 𝔼(*f*^2^(Φ_*a*_)) and 𝔼(f ^2^(Φ_*A*_)) = 𝔼(*f*^1^(Φ_*a*_)). This condition also ensures that the average of two alleles’ mean fitnesses, which we denote 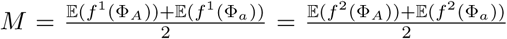, is the same in both environments. The function *f*^2^ is defined as a reflection of *f*^1^ around M: *f*^2^(x)= 2M − *f*^1^(x). As a result, the variance in fitness of allele a with randomly drawn phenotype is the same in both environments: Var(*f*^1^(Φ_*a*_))=Var(*f*^2^(Φ_*a*_)) =Var(Φ_*a*_). The symmetry conditions we have imposed on phenotypic means allow us to focus our analysis on the effects of phenotypic variation alone. These fitness functions describe a general model in which each genotype has a preferred environment, but allele *a* can express a range of phenotypes. For our analytical treatment we assume that the *a* allele has access to only two phenotypic values, as depicted in **Figure 1B**; but **Figure 6** shows similar results in simulations of a model with more than two phenotypic states associated with allele *a*.

**Figure 1:**
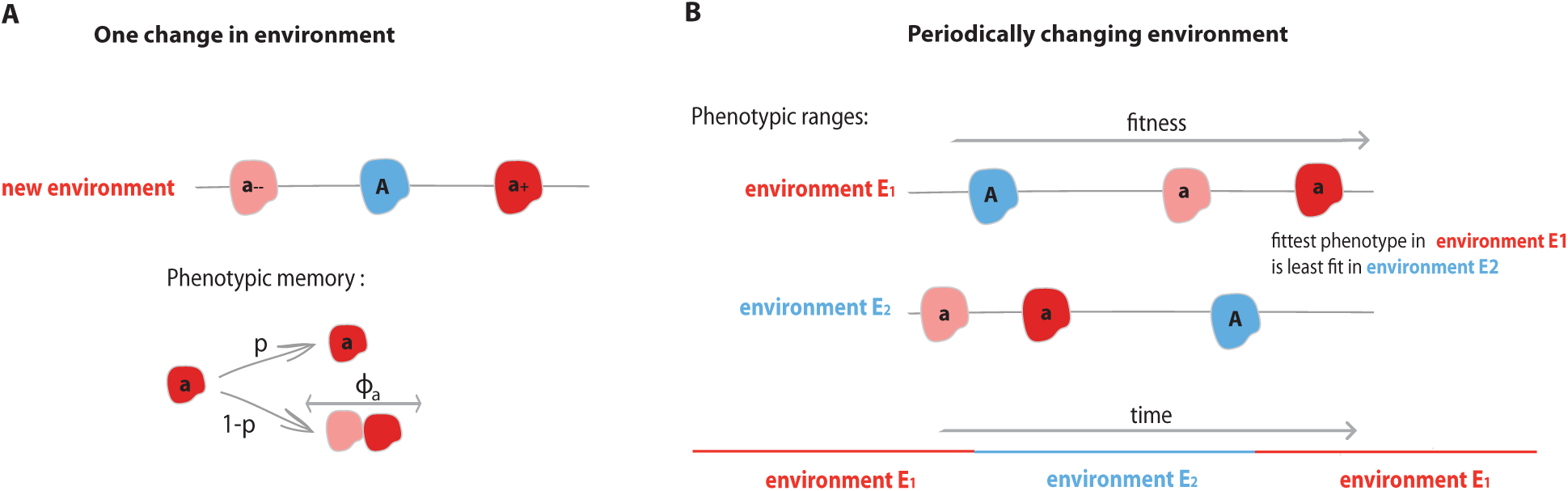
Illustration of the two versions of the model. We study both one abrupt change in the environment (**Panel A**), as well as periodic changes in selective pressure (**Panel B**). In both cases, when an individual *a* gives birth, with probability *p*, the probability of phenotypic memory, its offspring inherits the phenotype of its parent, and with probability 1 − *p*, the offspring’s phenotype is resampled from the phenotypic distribution.

We study the possible long-term advantage of heritable phenotypic variability by analyzing how the introduction of the *a* allele into an otherwise non-variable population (*A*) changes the population’s probability of rescue or mean times to extinction. We quantify how the probability of rescue or the time to extinction depends on environmental factors, such as the environmental period 2*n*, on demographic factors, such as the carrying capacity *K*, and on molecular factors, such as the variance in phenotypes that can be expressed by allele *a*, Var(Φ_*a*_), and the degree of phenotypic memory, *p*. We derive analytic approximations in the case of one environmental change, and we determine the mean times to extinction in changing environments by Monte Carlo simulations (Gillespie, 1976), using an ensemble of at least 10,000 replicate populations.

We simulate the birth-death process in continuous time as follows. We sample the waiting time for an event from an exponential distribution with rate parameter equal to the sum of all possible rates beginning at time zero; we then randomly assign a specific event according to the relative probabilities of occurrence of each event type (birth or death events) and update the population status, time, and all event rates. If the event implemented is a birth we then determine the phenotypic state of the offspring as follows. If the individual chosen to reproduce has genotype *A*, then the phenotypic state of the offspring always equals its parent’s (fixed) phenotypic value. For a reproducing individual with the a allele, however, there exists a probability of phenotypic memory, denoted by the parameter *p*, between parent and offspring: with probability *p* the offspring retains the phenotypic state of its parent, and with probability 1 − *p* the offspring’s phenotype is redrawn independently from the random variable Φ_*a*_. Thus, individuals of type *a* can express a range of phenotypic values, and their phenotype is partly heritable between generations (provided *p* > 0). In the case of periodic environments, we implement environmental changes (and re-calculate event rates) at deterministic times: *n*, 2*n*, 3*n*, etc. Time is measured in units of an individual’s expected lifetime – that is, the death rate is set to unity for all individuals in all simulations.

## Results

### Evolutionary rescue from a novel mutation

We study the probability of evolutionary rescue after a single, abrupt environmental change that would lead to population extinction of the wild-type allele *A*. We assume that there is a constant rate of mutation from the *A* allele towards the *a* allele, *μ*. We start by obtaining analytic approximations for the density-dependent model and then discuss the density-independent case. Intuitively, the density-independent model applies in biological situations where the population is very far from carrying capacity K (such that the term 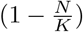 is approximately 1), while the density-dependent case applies for populations large enough at the time of environmental change that 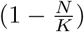 cannot be ignored.

### Density-dependent births

The probability of establishment of the new mutant (and therefore the probability of rescue) depends critically on the mutation rate to the *a* allele, and whether the first *a* individual is initially introduced with its beneficial or its deleterious phenotype – that is, whether its birth rate is initially larger or smaller than its death rate. We first study the case where the A allele can only mutate to produce an a allele with the beneficial phenotype, denoted by Φ_*a*_+. Biologically, this could be due to the presence or absence of an epigenetic marker that makes the deleterious Φ_*a*_phenotype inaccessible directly from the A allele. The population will be rescued, by definition, if the a lineage manages to become established (Uecker and Hermisson, 2011). As shown in **Figure 2A**, the chance of evolutionary rescue increases monotonically with the strength of phenotypic memory, p. This result makes intuitive sense: high-fitness variants of the a allele are preferentially transmitted to the next generation, and greater phenotypic memory p increases their propensity to maintain the high-fitness phenotype and become established in the population. Moreover, the probability of rescue is uniformly greater when the a allele can express a greater diversity of phenotypes, i.e. for Var(Φ_*a*_) large (**Figure 2A**), because the larger variance is associated with a greater fitness for the Φ_*a*_+ phenotype. In summary, when the variable allele is introduced with a beneficial phenotype, rescue is facilitated by increased phenotypic memory and by increased phenotypic variance of the phenotypically-variable allele.

**Figure 2:**
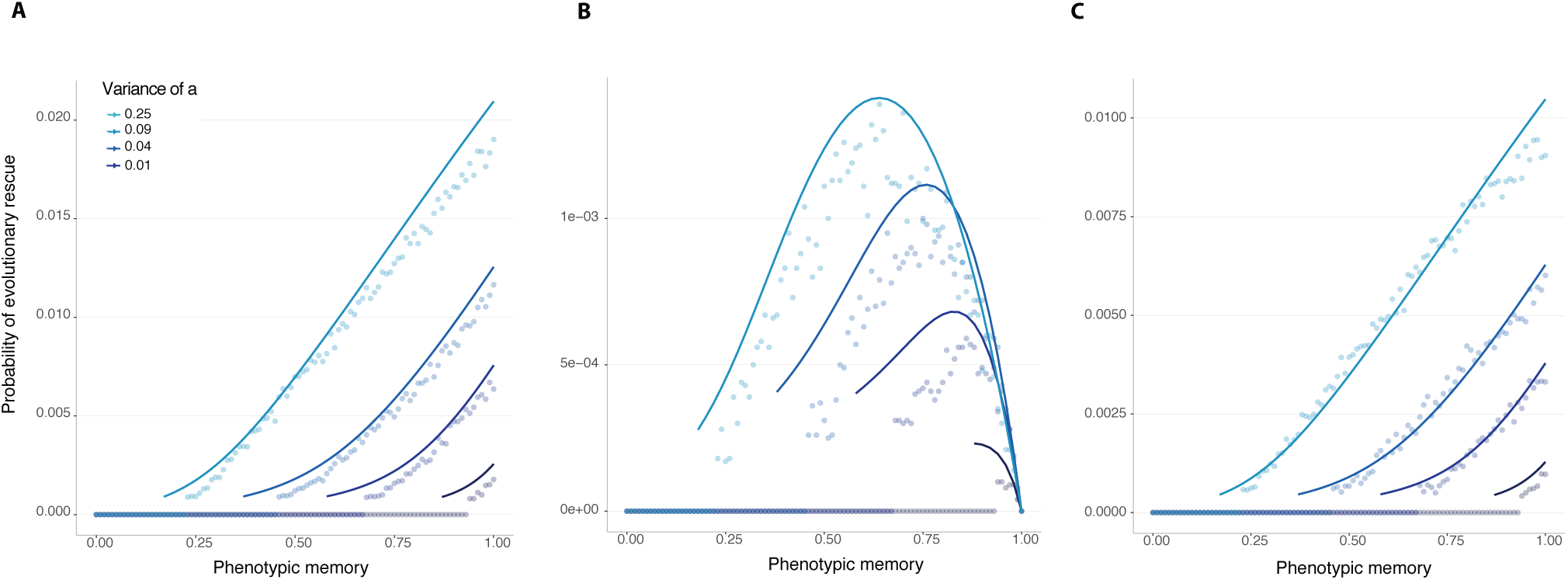
Probability of evolutionary rescue from a new mutation, with density-dependent births. The lines represent the analytical approximations. The dots represent the ensemble average across 10,000 replicate Monte Carlo simulations. Here, *K* = 5000, *N*_0_ = 1000, *N*_0_*μ* = 0.01 and 𝔼(Φ_*A*_)= 𝔼(Φ_*a*_)= 0.95. **Panel A**: Probability of rescue when A can only initially access the a_+_ phenotype. **Panel B**: Probability of rescue when A can only initially access the a_*-*_ phenotype. **Panel C**: Probability of rescue when A can only initially access both a phenotypes.

When the *a* allele is introduced with a deleterious phenotype Φ_*a −*_, evolutionary rescue can still occur, because the phenotype of type-a individuals may change between generations. In this case, **Figure 2B** shows that the probability of evolutionary rescue depends non-monotonically on the strength of phenotypic memory p. There is simple intuition for this result as well, which is informed by our mathematical analysis below. Intuitively, the probability of establishment in this case is the product of the probability that some a-type individual produces an offspring with the beneficial phenotype, 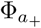, before the a-lineage is lost, times the probability of establishment associated with such an individual with phenotype 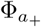. Therefore, rescue is facilitated as the strength of phenotypic memory increases above zero (this effect is driven by the increase in the probability of rescue once an individual of phenotype 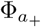 arises); but as the phenotypic memory increases further, towards one, the probability of rescue is reduced, because the entire a lineage will likely go extinct before producing any individual with a beneficial phenotype. Or put differently, the a lineage needs sufficient variability to produce the correct phenotype, but not too much to avoid losing it after it has been produced.

To provide a clear analysis of the intuitions described above, we first derive the probability of rescue, ℙ_*r*_(a_+_), when there is recurrent mutation towards the beneficial phenotype 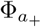. To do so, we first compute an effective selection coefficient of the entire, phenotypically variable a lineage, by assuming that the two phenotypes within the a lineage quickly reach mutation-selection balance. Given phenotypic “mutation” rate 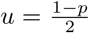 between the two phenotypes, at equilibrium, the frequency of phenotype 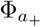 within the a-lineage is given by f_*a*_+:

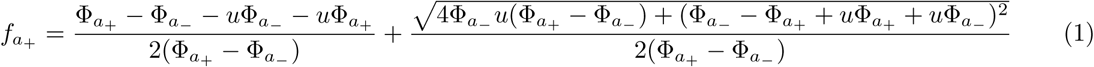

We then compute the effective birth rate of the a lineage

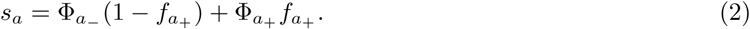

This can be rewritten as a function of the mean 𝔼(Φ_*a*_) and the standard deviation 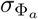 of the two alleles as:

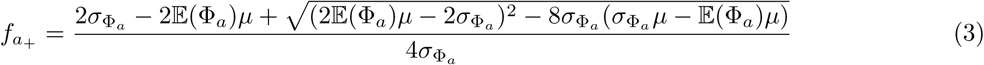

and

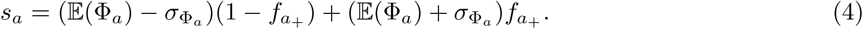

The effective birth rate s_*a*_ determines the equilibrium population size for the adapted population, should adaptation and rescue occur: 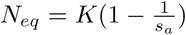, assuming s_*a*_ > 1. When the effective birth rate *s*_*a*_ of the a lineage exceeds its death rate (unity), the probability of establishment of this new mutation arising at time τ in the population is given by

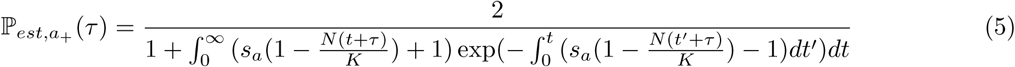

following Uecker and Hermisson (2011) and Wilson et al. (2017). Assuming that the mutant lineages have independent probabilities of establishment, and thus neglecting the mutant population size, *N*(*t*) can be approximated by the *A* allele population size, which, due to the density-dependence in birth rate for the wild-type A, has the form

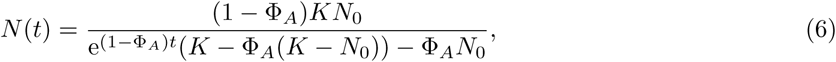

where *N*_0_ = *N*(*t* = 0).

We can now derive the probability of evolutionary rescue from at least one adaptive mutant by modeling mutant establishments using a time-inhomogeneous Poisson process with intensity function *μN*(*t*)*P*_*est,a*_+. The probability of evolutionary rescue then becomes one minus the probability that no mutants establish:

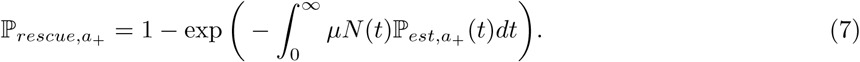

This analytic approximation is shown in **Figure 2A** alongside the results of Monte Carlo simulations. For comparison to simulation, we defined rescue as the population reaching 99% of the equilibrium population size of the adapted population N_*eq*_. We also count any cases in which N_*eq*_ was smaller than 100 as extinction (because population sizes so small can easily fluctuate to extinction).

Conversely, when the a mutation is introduced with its deleterious phenotypic state, Φ_*a*_, we derive an approximation for the probability of establishment ℙ_*est,a* −_(t) as the probability of at least one phenotypic mutation to 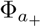 before the loss of the a allele, multiplied by 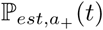. In other words, if we let η denote the event that there is at least one phenotypic mutation within the a-lineage before its loss, then we will approximate the probability of establishment as

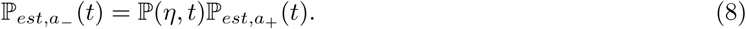

We need only derive an expression for *P*(*η*, *t*). The probability that no mutation occurs before its loss is

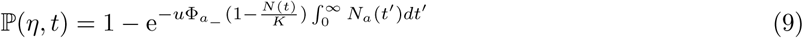

according to mutation viewed as a time-inhomogeneous Poisson process with intensity function 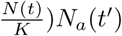. Here N_*a*_(t^′^) represents the decline in the a population from a single individual with negative expected growth rate:

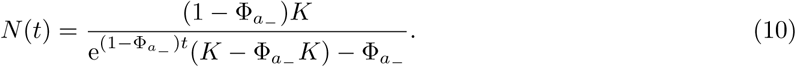

If we make the approximation that the switch to the beneficial a phenotype is instantaneous, then the analytic expression for the probability of rescue becomes

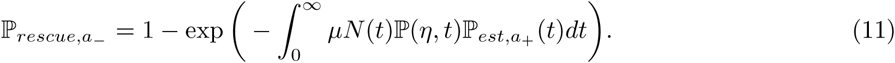

This analytical approximation is shown in **Figure 2B**, alongside the results of an ensemble of Monte Carlo simulations. In **Supplementary Figure S1** we show the analytic approximation along with evolutionary simulations for a range of rates of mutation.

Finally, we can approximate the probability of rescue when novel mutations to the a allele are introduced with random phenotype 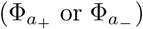 with equal probability (**Figure 2C**). Although there are several ways to make an analytic approximation, the simplest is simply to note that the probability of rescue from mutation to 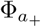 is larger by an order of magnitude than the probability of rescue from mutation to Φ_*a*_. Therefore we can approximate the overall probability of rescue with random initial phenotype as the probability of rescue from mutation to 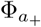 using mutation rate 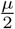. Using equation (7), this gives

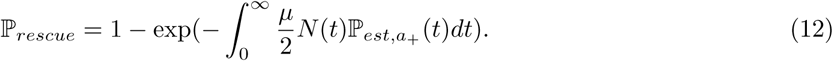

### Density-independent births

The analytic results above can be simplified greatly when the population is far from carrying capacity *K* at the time of the environmental shift. In this regime we can effectively assume that both the birth rates and the death rates are density-independent. In this case we can again compute the effective birth rate for the a lineage, s_*a*_. If the effective birth rate s_*a*_ is lower than the death rate, then we say there is zero probability of rescue. If the effective birth rate *s*_*a*_ exceeds unity, then equation (7) simplifies to

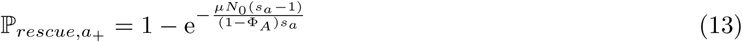

taking into account that, in this regime, the population size can be deterministically approximated by *N*(*t*)= *N*_0_ exp(*-*(1 Φ_*A*_)t. Similarly, the probability of rescue from recurrent mutation to Φ_*a* −_becomes

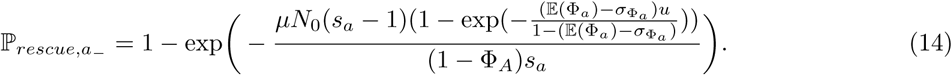

These two analytic approximations are shown in **Figure 3A** and **B**, alongside the results of Monte Carlo simulations. For comparison to simulations in the density-independent case we defined rescue as the population reaching size *N* = *K*.

As expected, the density-independent approximation for the rescue probability is similar to the full density-dependent treatment for systems starting far from carrying capacity (**Supplementary Figure S2**).

For the sake of simplicity the analysis that follows (cases of rescue from standing genetic variation or rescue from a resistance mutation) assumes that birth and death rates are density-independent.

### Evolutionary rescue from standing genetic variation

Evolutionary rescue may alternatively occur from an adaptive variant that exists, at low frequency, at the time of the environmental shift that dooms the wild-type genotype to extinction. To study this we assume that at time *t* =0 the population already contains a proportion *q* of *a* alleles, and we study the probability of rescue from this standing genetic variation (**Figure 4**). In this case the *a* lineage is assumed to be at mutation selection balance between its two phenotypic forms at the time of the environmental change *t* = 0. The mutation selection balance is computed using equation (1) and the probability of rescue becomes

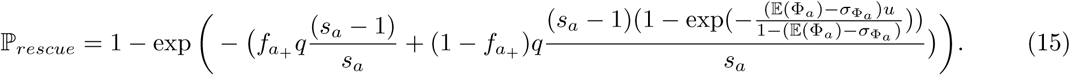

The result of this approximation, together with Monte-Carlo simulations are shown in **Figure 4**. As the phenotypic memory *p* increases, the phenotypic mutation towards *a*_*-*_ decreases, and therefore the fraction of *a*_*-*_ phenotypes initially present in the population decreases. Since the probabilities of establishment for the deleterious a_*-*_ phenotypes are smaller than the ones for the a_+_ phenotypes, the initial number of a_+_ phenotypes is the main driver of the evolutionary dynamics.

**Figure 3:**
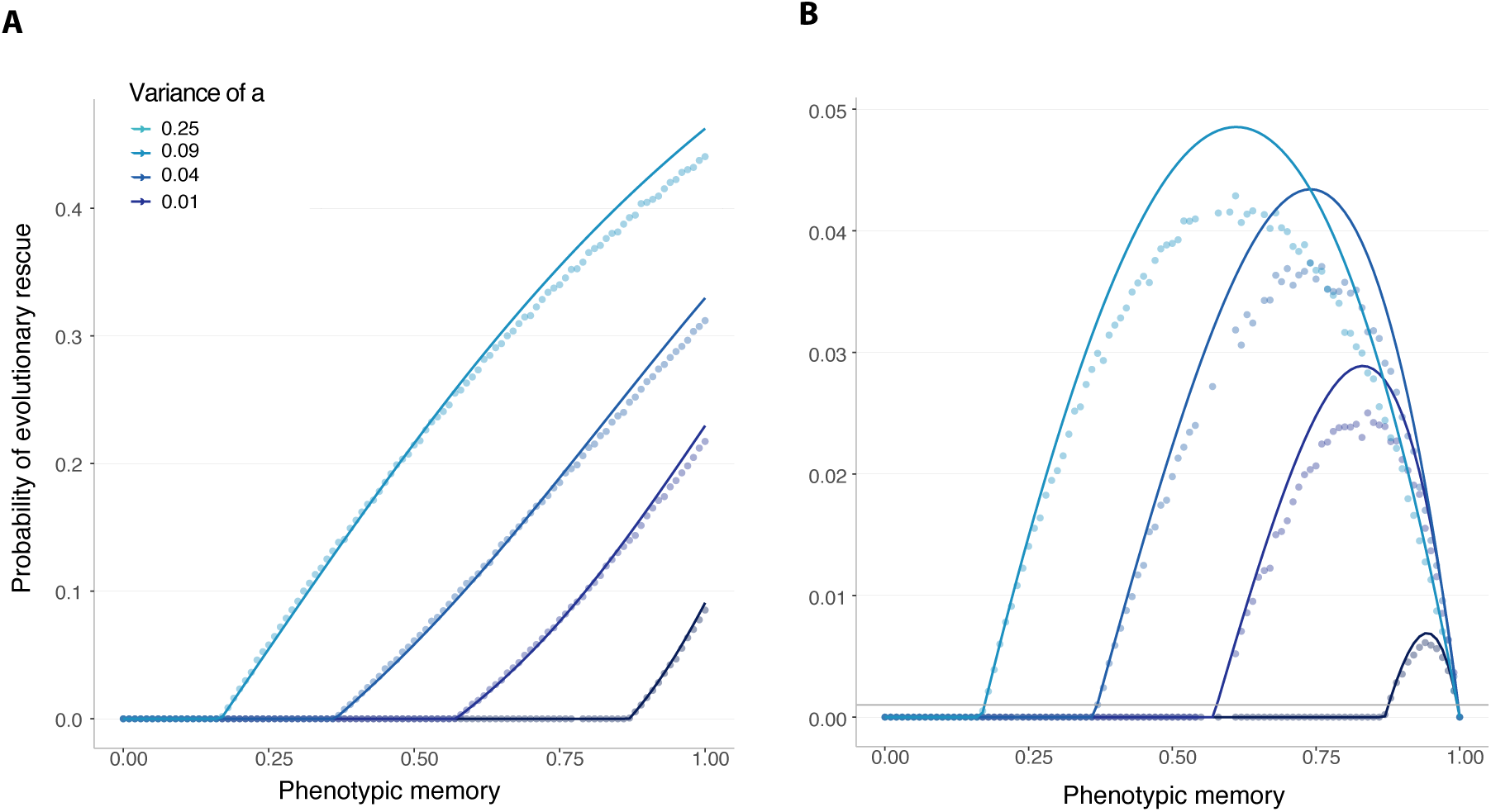
Probability of evolutionary rescue from a new mutation, density-independent birth. The lines represent the analytical approximations. The dots represent the ensemble average across 10,000 replicate Monte Carlo simulations. Here, *N*_0_ = 1000, *N*_0_*μ* = 0.1 and 𝔼(Φ_*A*_) = 𝔼(Φ_*a*_) = 0.95. **Panel A**: Probability of rescue when A can only initially access the a_+_ phenotype. **Panel B**: Probability of rescue when A can only initially access the a_*-*_ phenotype.

**Evolutionary rescue from a novel resistance mutation**

We adapted our model slightly to study the idea that phenotypic variability might allow the population to persist until a more permanent genetic change, such as a resistance mutation, can establish. To study this we introduced a second, linked locus, *M*, into our model, at which a resistance mutation can appear at a rate smaller than the mutation rate at the *A*/*a* locus. We compare the probability of evolutionary rescue due to a resistance mutation in populations fixed for the non-variable allele *A* versus populations that allow the phenotypically variable a allele. We find that, indeed, phenotypically heterogeneous populations have an advantage over populations composed of individuals of fixed phenotype, regardless of the strength of mutation at the resistance locus, *M* (**Supplementary Figures S3** and **S4**).

**Figure 4:**
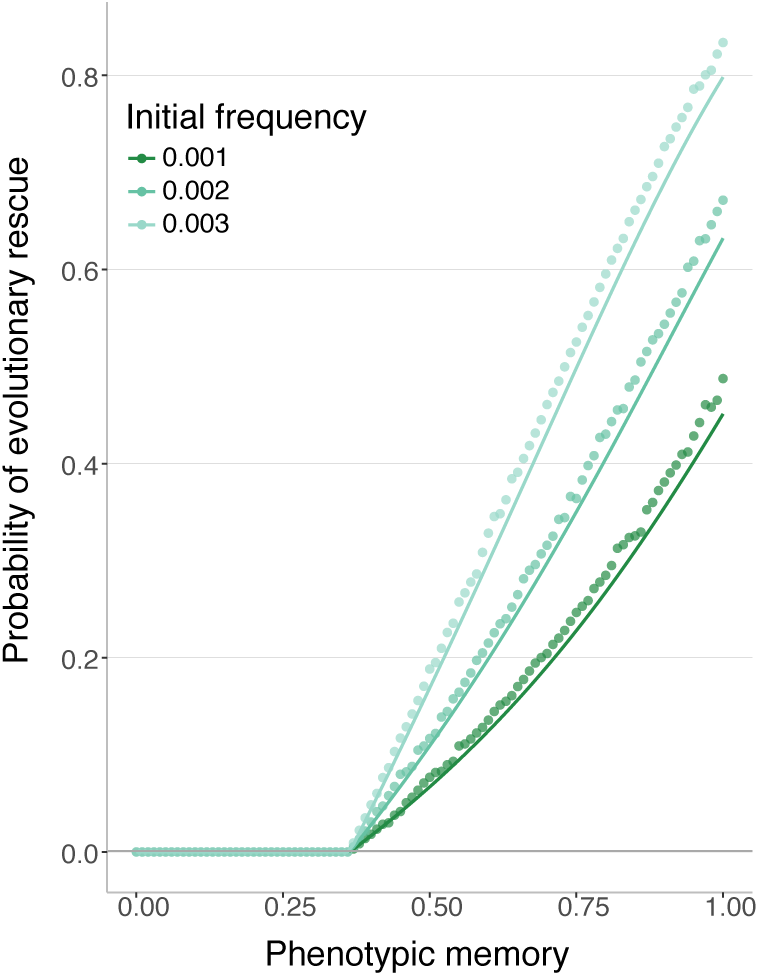
Probability of evolutionary rescue from standing genetic variation, density-independent birth. The lines represent the analytical approximations. The dots represent the ensemble average across 10,000 replicate Monte Carlo simulations. Here, *N*_0_ = 2500, 𝔼(Φ_*A*_) = 𝔼(Φ_*a*_) = 0.95 and Var(Φ_*a*_)= 0.09.

### Population persistence in periodically changing environments

Phenotypic variability and phenotypic memory also influence population persistence in periodically changing environments, in addition to the case of a single environmental change that has been the subject of most work on evolutionary rescue. With a periodically changing environment, the question of persistence is conveniently framed in terms of the mean time before population extinction. Even in this more complicated setting we once again observe a non-monotonic effect of phenotypic memory, *p*, on population persistence: populations go extinct quickly for either small *p* or large *p*, whereas intermediate amounts of phenotypic memory can promote persistence for long periods of time.

**Figure 5A** shows the mean time to extinction as a function of the phenotypic memory, for a range of environmental periods *n*. In all these cases, a population comprised of only the non-variable wild-type allele A goes extinct quickly (cf. **Supplementary Figure S5**). But populations initiated with a single copy of the phenotypically-variable allele a have the potential to persist for very long times, especially for intermediate values of the phenotypic memory parameter, *p*.

In our model of periodic environments, faster environmental changes are correlated with longer popu-lation persistence, even in the case of a phenotypically invariant wild-type population (**Supplementary Figure S5**). This occurs because long stretches of the environmental regime deleterious to the *A* allele lead to population declines that the beneficial periods cannot replenish. It is particularly in faster-changing environments that phenotypic memory in a phenotypically-variable allele a provides the largest advantage for population persistence – because it helps the high-fitness realizations of a allele remain high-fitness, which is essential for population persistence through environmental epochs deleterious to the wild-type *A* allele. The non-monotonic dependence of persistence time on phenotypic memory also makes intuitive sense. On the one hand, it is beneficial for the a allele to have some phenotypic memory within each environment (*E*_1_ or *E*_2_), as this helps the high-fitness realizations of the allele with little effect on its low-fitness realizations. On the other hand, too much phenotypic memory is detrimental in the long-run, because once the environment shifts, the a lineage will be “stuck” with a deleterious phenotype. Moreover, regardless of the phenotypic memory, the duration of persistence always increases with the variance in phenotypes that a can express, Var(Φ_*a*_) (**Figure 5B**) – that is, the population can persist longer when the phenotypically-variable allele has access to a larger phenotypic range.

**Figure 5:**
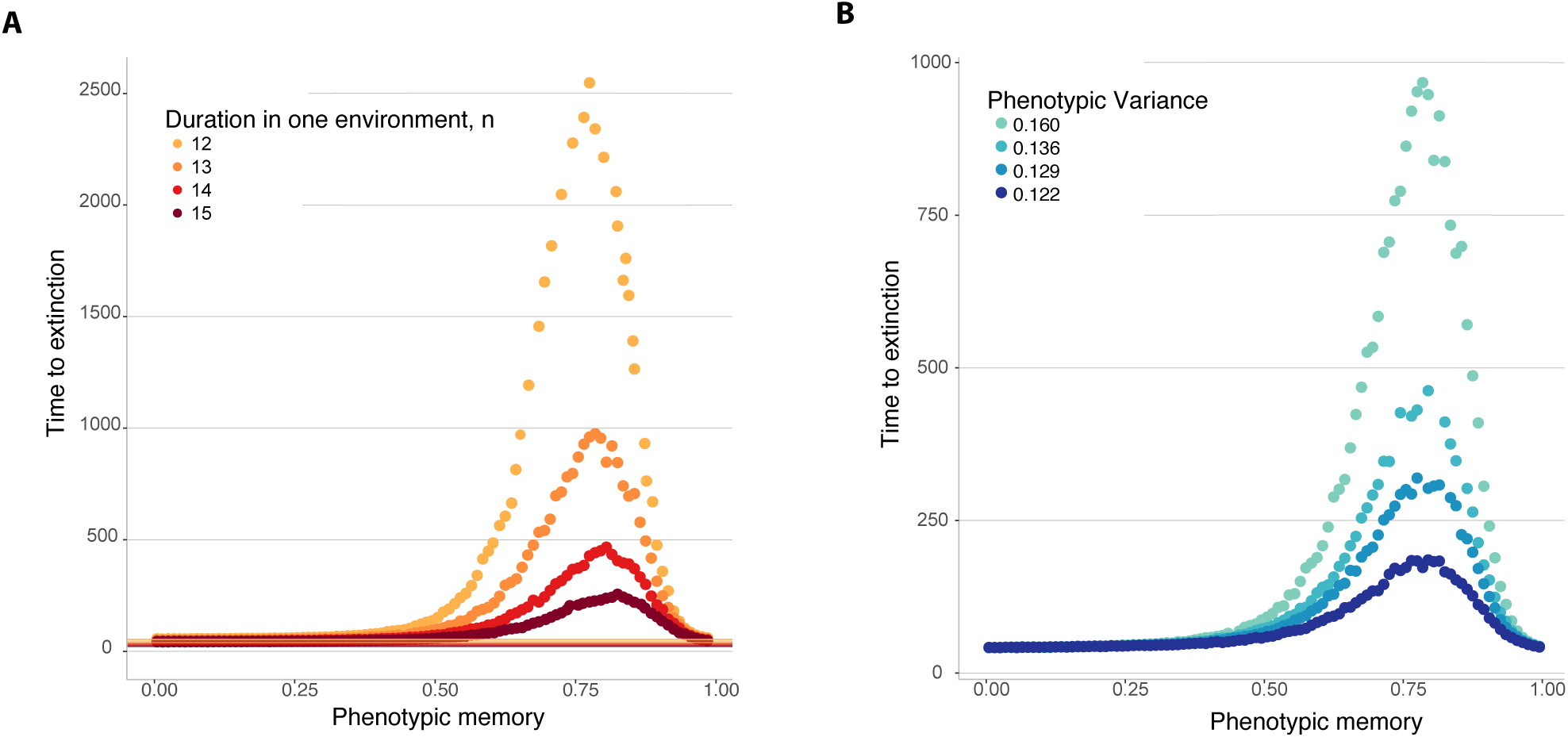
Population persistence in periodically changing environments. The dots represent the mean time before population extinction across 10,000 replicate simulations. All populations are initiated at half carrying capacity, *N* = 500, with a carrying capacity *K* = 10000, with a single mutant genotype a drawn with a random phenotype introduced into a random one of the two different environments. The two environments then cycle deterministically with each environmental epoch lasting n time units, where time is measured in units of the expected lifespan of an individual. **Panel A**: Mean time until population extinction for a range of different environmental periods, with *f*^1^(Φ_*A*_) = 0.5, *f*^2^(Φ_*A*_) = 1.5 and Var(Φ_*a*_) = 0.16. The lines, each corresponding to a different value of n, show the mean time to extinction of a population comprised of only wild-type A alleles. **Panel B**: Mean time until population extinction for different amounts of phenotypic variability, Var(Φ_*a*_), with a fixed environmental duration of *N* = 13 units, *f*^1^(Φ_*A*_) = 0.5, *f*^2^(Φ_*A*_)= 1.5.

### Multiple phenotypes

We also explore the probability of rescue and time to extinction for populations in which the a allele has access to more than two different phenotypes. In **Figure 6A** and **B** we show by simulation that, when the a allele has access to four or six different phenotypes, as long as the phenotypic variance of the a allele is kept constant, the rescue probabilities are virtually the same as in the case of a phenotypic random variable with two point masses. The same holds for the time to extinction in periodically changing environments (**Figure 6C**). In **Supplementary Figure S6** we vary the number of phenotypes available to the a lineage while keeping its phenotypic range constant (increasing the number of phenotypes available with a constant phenotypic range decreases the phenotypic variance). As we might expect, for a lineages with the same phenotypic range (and therefore the same maximum growth rate), the probability of evolutionary rescue decreases in accordance with the accompanying decrease in the phenotypic variance.

**Figure 6:**
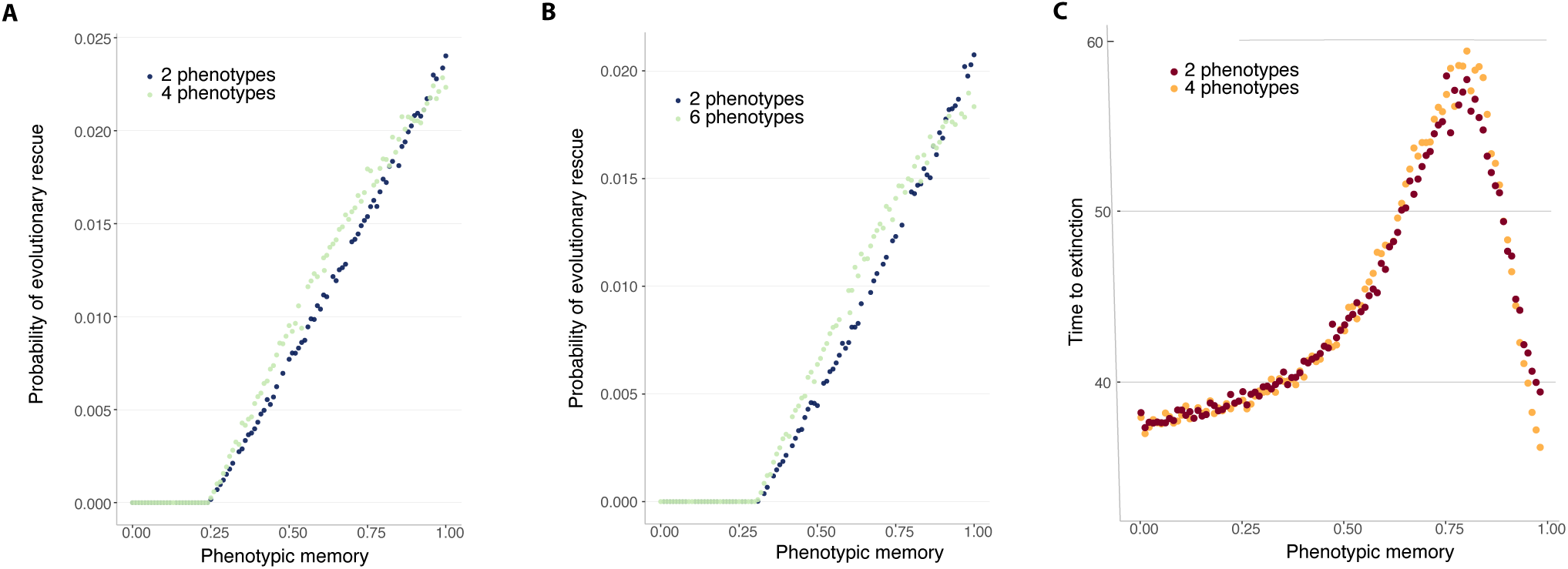
Probability of evolutionary rescue and time to extinction when the a allele has access to more than two phenotypes. Panel A: Here, *N*_0_ = 1000, *N*_0_*μ* = 0.01 and 𝔼(Φ_*A*_) = 𝔼(Φ_*a*_) = 0.95, with Var(Φ_*a*_)=0.156. **Panel B:** Parameters are the same as in **Panel A**, with Var(Φ_*a*_)=0.11. **Panel C:** Parameters are the same as in **Figure 5B** with Var(Φ_*a*_)=0.076.

## Discussion

Evolutionary adaptation occurring on the same timescale as demographic dynamics can have profound effects on the persistence of a population. The theory of evolutionary rescue provides a conceptual framework that links demography and evolution in finite populations of variable size (Lindsey et al., 2013; Uecker and Hermisson, 2015; Wilson et al., 2017; Uecker et al., 2014; Orr and Unckless, 2008, 2014; Alexander et al., 2014; Bertram et al., 2016). Populations experiencing a sudden change in selection pressures or, frequent and unpredictable environmental variation, may either genetically adapt or be unable to recover. These populations have a limited window of opportunity for individuals with phenotypic solutions advantageous in the novel environment to appear and establish. Genetic adaptation after abrupt environmental changes can prevent extinction under several different demographic scenarios, but such a mechanism of rescue is inherently limited by the genomic mutation rate.

Transient and variable phenotypes, which can be mediated by rapid transitions in the epigenome, may provide an additional, selectable layer of traits that enable populations to persist until the appearance of more permanent strategies, such as genetic resistance. In systems ranging from bacterial infections, to latency in viral populations, or cellular neoplastic development, this form of epigenetic, partly heritable phenotypic heterogeneity has been shown to facilitate adaptation and population persistence under changing selection pressures (Seger and Brockman, 1987; Acar et al., 2008; Veening et al., 2008). These responses via partially heritable phenotypic variability can occur on faster time scales than genetic responses, and they may play a critical role in the path towards long-term resistance eventually reinforced by genetic changes. Indeed, here we explored the fate of populations waiting for a resistance mutation, and we found that that phenotypically variable populations have a higher probability of rescue via resistance than less phenotypically variable populations.

The goal of this study has been to develop a population-genetic model to quantify the probability of evolutionary rescue and mean times to extinction for populations that can access an allele with increased, partly heritable phenotypic variation. We have studied the problem of evolutionary rescue both from a novel mutation, or for pre-existing standing variants. Our analytical results are restricted to the case in which the phenotypically-variable lineage can access two different phenotypes, but we have also seen that the results hold in simulation for phenotypic random variables with more than two phenotypic states. The model with exactly two phenotypic states can be easily reframed as a model of stochastic phenotypic switching. Here the switching rate can be written as 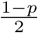, where *p* is the phenotypic memory described in this paper. This reframing helps place some of our results in the context of the larger literature on phenotypic bet-hedging that has been developed both for infinite and finite populations and for environments fluctuating through time (Carja and Plotkin, 2017; Carja et al., 2014a; Kussell et al., 2005; Balaban et al., 2004; Salathe et al., 2009).

It is widely recognized that phenotypic variation can serve as a response to environmental change(Botero et al., 2015; Tufto, 2015). Previous theoretical studies on the adaptive benefit of stochastic phenotypic variability have mostly analyzed the evolution of rates of phenotypic switching by modifier loci, and mostly in infinite populations (Carja et al., 2014a; Salathe et al., 2009). These studies have found that phenotypic switching rates should evolve in tune with the correlation between the environments of parent and offspring: when the environment fluctuates periodically between two states with different optimal phenotypes, the uninvadvable switching rate (the ESS) between phenotypic states will evolve to approximately 1/n, where n is the number of generations between environmental changes (Ishii et al., 1989; Salathe et al., 2009). Further studies generalized these results to include both environmental and spatial fluctuations in selection (Carja et al., 2014a,b). We have previously explored the dynamics of phenotypic switching rates in finite populations of fixed size, where we found that the fixation probability of phenotypically-variable alleles depends nonmonotonically on the probability of phenotypic inheritance between parent and offspring: probabilities of fixation are maximized at intermediate rates of phenotypic switching in fluctuating environments (Carja and Plotkin, 2017). Here we find similar results for phenotypic switching rates in populations of changing size that experience periodic fluctuations in selective pressure (Figures 5 and 6): there exists an intermediate phenotypic switching rate that maximizes population persistance. This intermediate phenotypic mutation rate holds not just for phenotypic distributions with two states, but also when the phenotypically-variable lineage has access to more than two phenotypes; and it is the phenotypic variance that drives the evolutionary dynamics.

Although our simple model neglects the myriad of mechanisms that give rise to phenotypic variability, it makes some general qualitative and quantitive predictions that should hold broadly and can inform the design of therapies in clinical contexts where population eradication is desired. Indeed, many clinical examples of therapy failure are now known to be caused by phenotypic heterogeneity, persistence or quiescent cellular states (Cohen et al., 2013; Deris et al., 2013).

By exploring the interplay between phenotypic memory and treatment period, our results suggest that two very different types of intervention will be effective. Both options stem from the fact that, unlike genetic changes, epigenetic or phenotypic changes are reversible. The existence of an intermediate phenotypic memory that maximizes the time to extinction suggests that effective interventions are treatments that disrupt the molecular memory to either extreme (*p* = 0 or *p* = 1). This would facilitate eradication by decreasing the chance of a phenotypically resistant type establishing before the population goes extinct. Of course, further detailed predictive models, specialized to particular populations and drug actions, are needed to formulate optimal therapies across the plethora of diseases where transient phenotypic variability drives treatment failure. But we expect the lessons learned from simple models, concerning the complex effect of phenotypic memory on persistence, to hold generally.

## Acknowledgments

We gratefully acknowledge support from the David and Lucile Packard Foundation, the U.S. Department of the Interior (D12AP00025), and the U.S. Army Research Office (W911NF-12-1-0552). This research was done using resources provided by the Open Science Grid, which is supported by the National Science Foundation award 1148698, and the U.S. Department of Energy’s Offce of Science.

---

**Figure S1:**
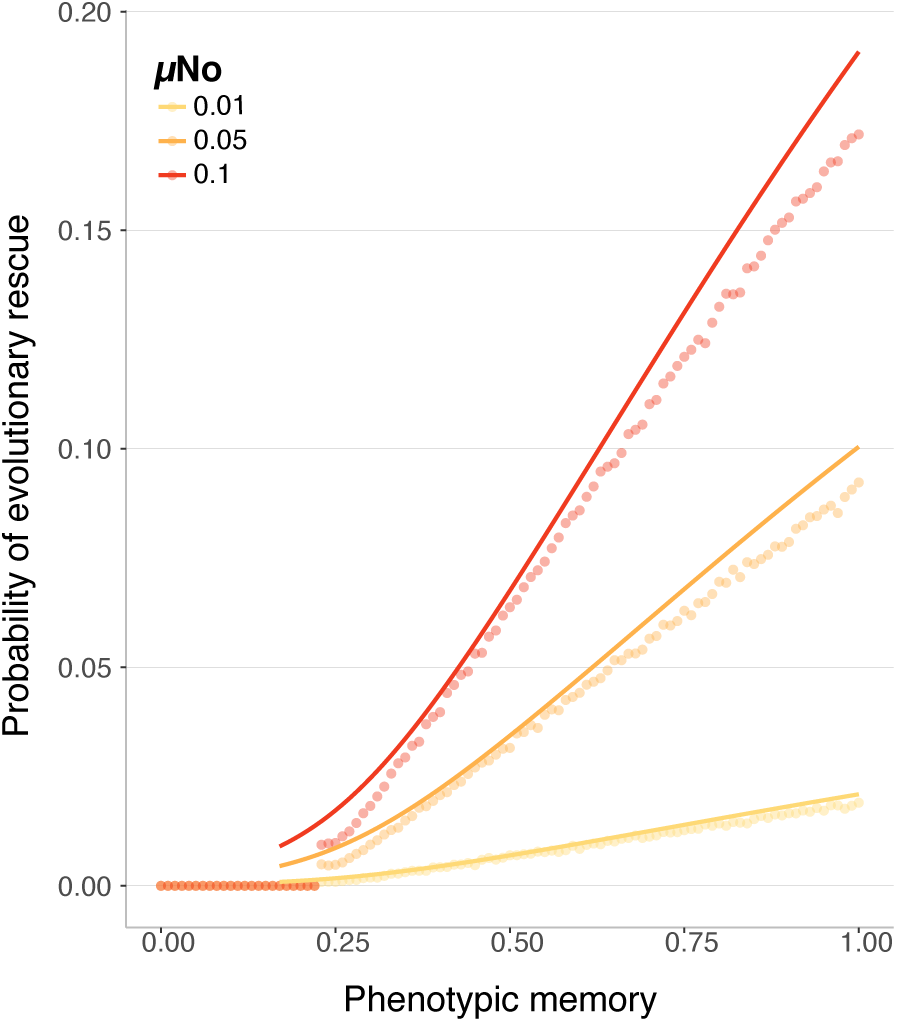
Probability of evolutionary rescue as a function of N*μ* from a new mutation density-dependent birth. The lines are analytic approximations, while the dots represent the ensemble average across 10,000 replicate Monte Carlo simulations. Here, the A allele can only mutate to the a_+_ phenotype, *K* = 5000, *N*_0_ = 1000, 𝔼(Φ_*A*_)= 𝔼(Φ_*a*_)= 0.95 and Var(Φ_*a*_)= 0.25.

**Figure S2:**
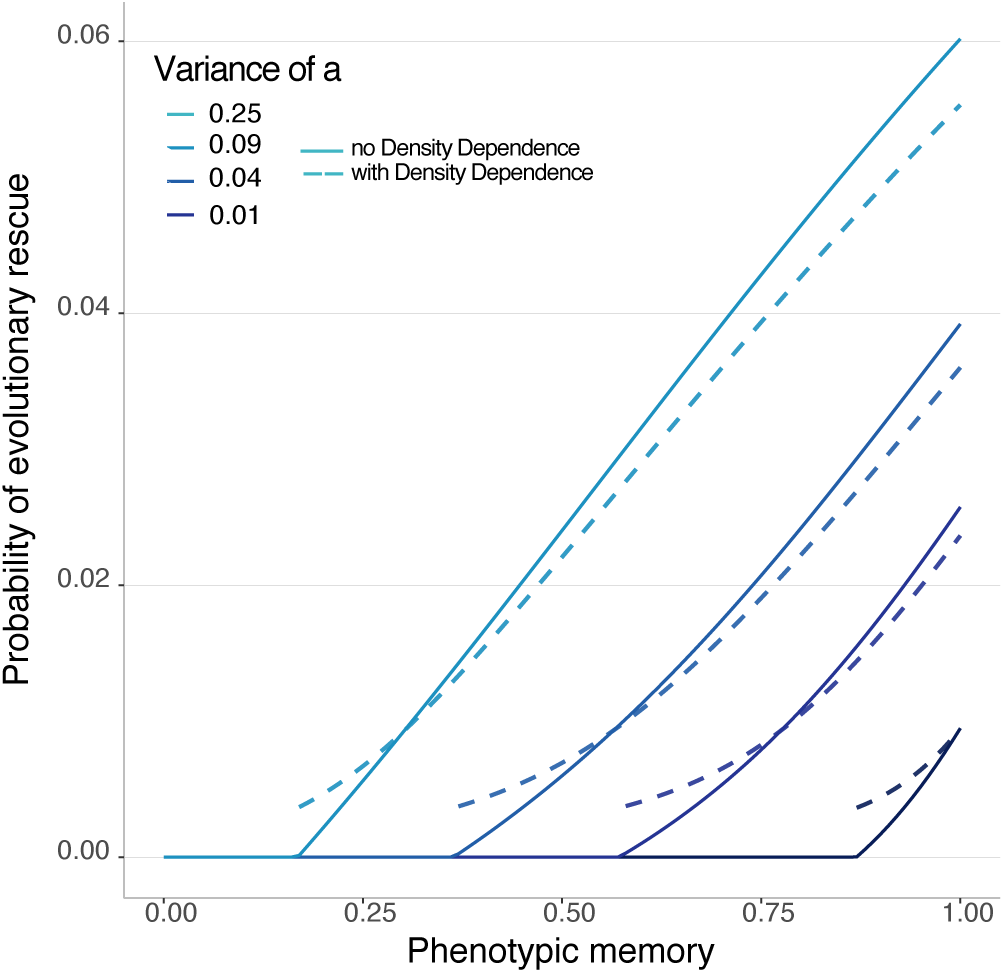
Comparison of density-independent and density-dependent analytic approximations, as 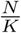 becomes small. Probability of evolutionary rescue from a new mutation. Here, *N*_0_ = 100, *K* = 1000000, *N*_0_*μ* = 0.01 and 𝔼(Φ_*A*_)= 𝔼(Φ_*a*_)= 0.95.

**Figure S3:**
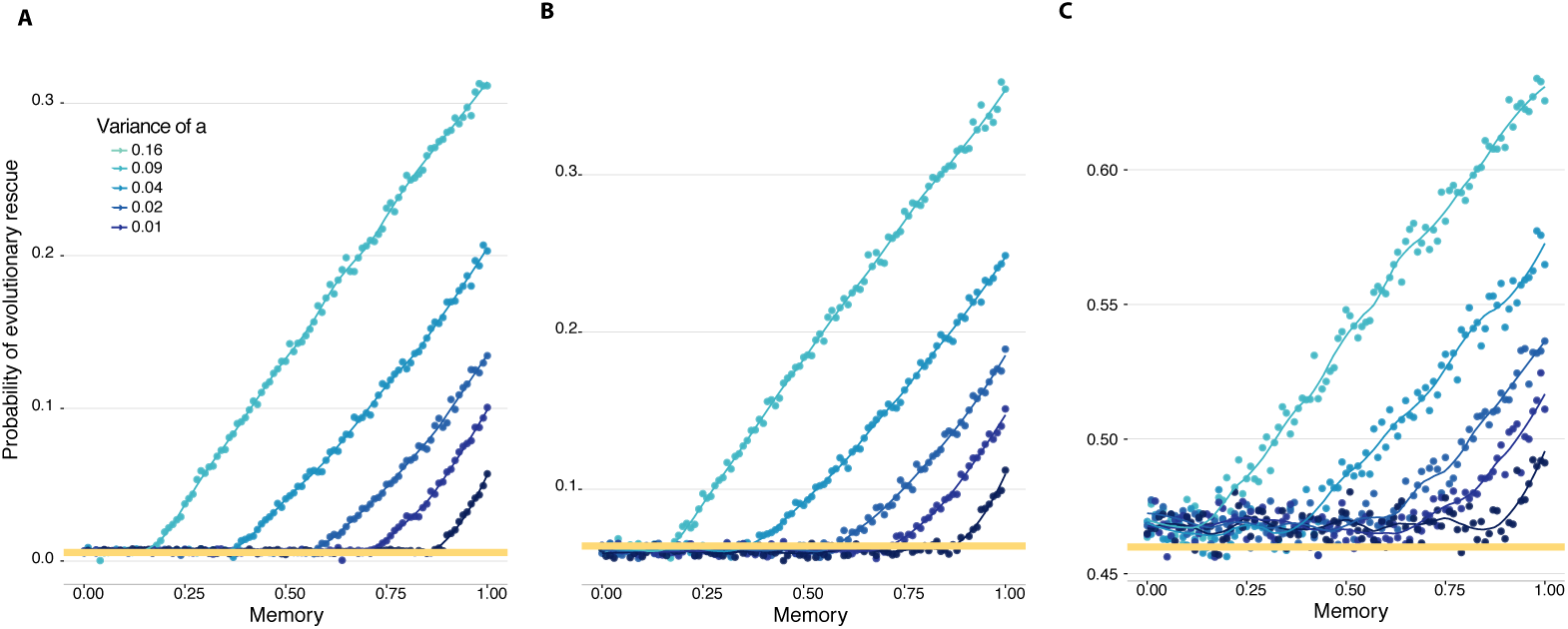
Probability of evolutionary rescue if there is a probability of genetic resistance, density-independent birth. The dots represent the ensemble average across 10,000 replicate Monte Carlo simulations. Here the mutation rate fromA to a, *μ*, is as presented in the panels. The mutation rate at the resistance locus, *μ*_*M*_, is assumed to be *μ*/100 and the growth rate of the resistance mutation is 1.5. **Panel A**: *μ* = 10^−5^. **Panel B**: *μ* = 10^−4^. **Panel C**: *μ* = 10^−3^. All populations are initiated at *N*_0_ = 2, 500, and 𝔼(Φ_*A*_)= 𝔼(Φ_*a*_)= 0.95. Yellow lines represent the probability of evolutionary rescue due to resistance for a population of fixed phenotype A.

**Figure S4:**
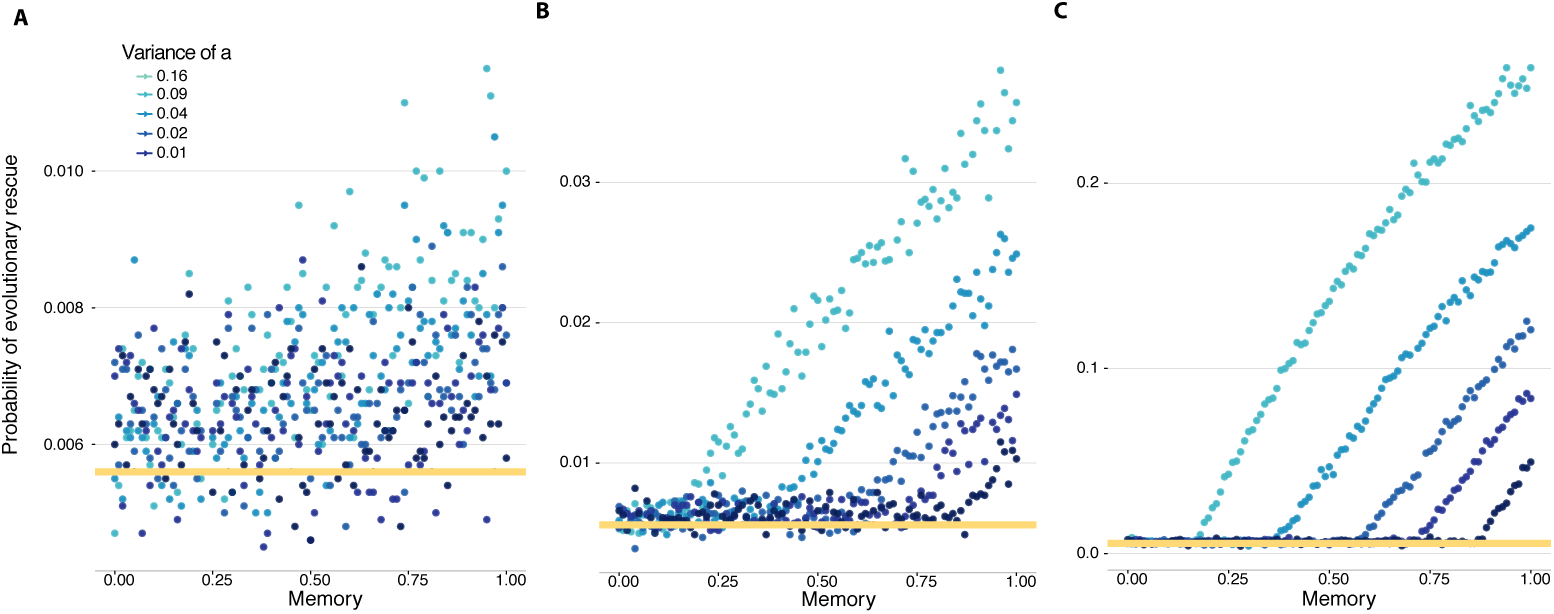
Probability of evolutionary rescue if there is a probability of genetic resistance, density-independent birth. The dots represent the ensemble average across 10,000 replicate Monte Carlo simulations. Here the mutation rate at the resistance locus is kept fixed at *μ*_*M*_ = 10^−7^ and the growth rate of the resistance mutation is 1.5. The different panels represent different rates of mutation, *μ* at the phenotypically-variable locus, from A to a. **Panel A**: *μ* = 10^−7^. **Panel B**: *μ* = 10^−6^. **Panel C**: *μ* = 10^−5^. All populations are initiated at *N*_0_ = 2, 500, and 𝔼(Φ_*A*_) = 𝔼(Φ_*a*_) = 0.95. Yellow lines represent the probability of evolutionary rescue due to resistance for a population of fixed phenotype A.

**Figure S5:**
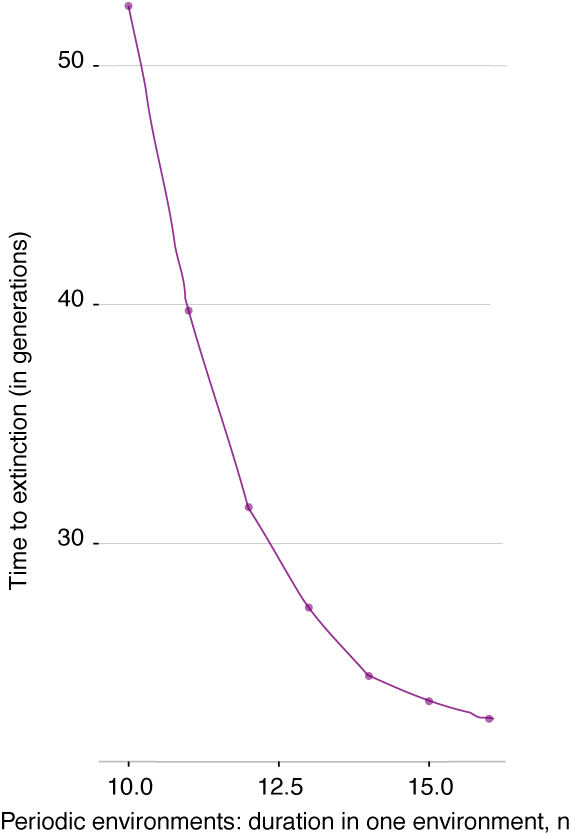
Mean time to population extinction for populations fixed on the wild-type. A. Here *f*^1^(Φ_*A*_)= 0.5, *f*^2^(Φ_*A*_)= 1.5 and the carrying capacity is *K* = 1, 000.

**Figure S6:**
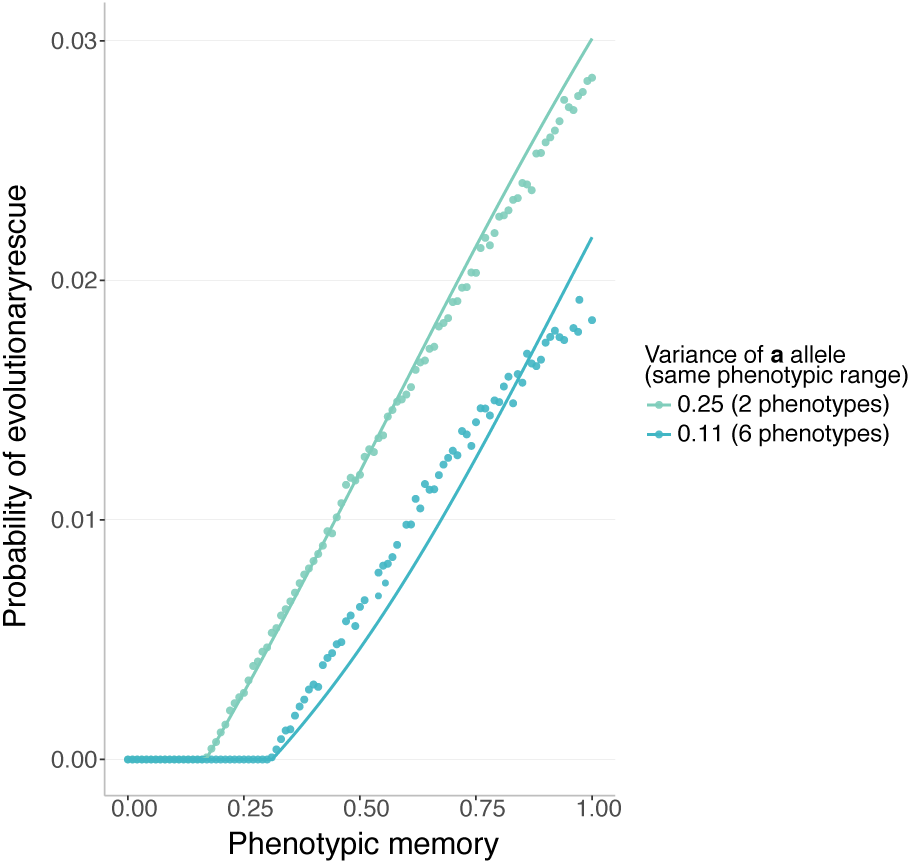
Probability of evolutionary rescue when the a allele has access to more than two phenotypes, same phenotypic range. Here, *N*_0_ = 1000, *N*_0_*μ* = 0.01 and 𝔼(Φ_*A*_)= 𝔼(Φ_*a*_)= 0.95, with the a allele with access to two or six phenotypes, such that its phenotypic range is constant (the maximum growth rate available to a is 1.45). The different numbers of phenotypes accessible to a determine the two different variances of the a allele phenotypes. The lines represent the analytic approximations computed using the right values of phenotypic variance. This figure shows that it is not the maximum growth rate available to a that drives evolutionary dynamics, and instead it is its variance.

